# The impacts of exogenous noise on stochastic disease dynamics

**DOI:** 10.64898/2026.09.09.750167

**Authors:** Daniel R. Higgins, Matt J. Keeling, Louise Dyson

## Abstract

Much of the literature and intuition associated with mathematical epidemiology is driven by deterministic models, which are a reasonable assumption when the population size is large. Stochastic models, especially individual based models, are however considered vital when dealing with small population sizes, especially at times of invasion or extinction. The overwhelming majority of these models (both deterministic and stochastic) assume that the underlying parameters are fixed (or follow a regular seasonal pattern). Here, we consider an analytic framework for dealing with randomly varying parameters through the use of stochastic differential equations - thereby capturing the action of external noisy processes such as weather. In particular, we focus on when the transmission rate, *β*, varies as the solution to a Cox-Ingersoll-Ross Model, such that *β* is gamma distributed with autocorrelation. We consider the impact of this parameter variation on a stochastic version of the Susceptible-Infected-Recovered model, and for this ‘double-stochastic’ model show through simulation and analytical results that exogenous noise increases the impact of stochasticity, potentially leading to more early extinctions, wider variations in the number of cases at equilibrium, but that early growth rate can be faster or slower depending on the precise parameters.

## 1. Introduction

Compartmental models are an extremely powerful method for modelling the dynamics of both outbreaks and endemic levels of infection. These models involve dividing a population into various classes (or compartments), such as *susceptible* or *infected*, forming a partition of the population. A system of ordinary differential equations (ODEs) can then be formulated to describe the rates of transition between these classes. These ordinary differential equation models can be very useful for modelling population-level processes [1, 2, 3, 4]. However, when the number of individuals in the susceptible or infected compartments is small, for example during invasion or eradication, the ODE assumptions break down and the individual nature of population needs to be considered [5, 6, 7]. Moreover, if one wishes to consider anything probabilistic in nature with relation to an outbreak, such as the extinction or invasion probability, a stochastic model is required.

One method for doing this, performed extensively in the literature, is to consider the epidemic as a continuous-time Markov chain (CTMC) whose state space corresponds to the integer number of individuals in each compartment. There exists a direct connection between the CTMC and the ODE system; the transition rates between states in the ODE are equivalent to the average corresponding rates in the CTMC. This means that as the population size is increased, the solution to the system of ODEs can be considered as the scaled limit of the CTMC [8]. In addition to introducing stochasticity to the model, a key difference is that the CTMC models *individuals* and so takes values in a discrete space; this allows for meaningful discussion surrounding extinction events and small population sizes.

The conversion of the ODE system to the CTMC system can be thought of as inducing *event-driven* randomness on the system while the expected rates do not change. That is, while the underlying rates of events for each individual do not change (or may change deterministically in time for forced systems [9]), the events are random in nature and thus individuals in a population can experience different outcomes due to randomness [10].

In addition to randomness of events, we can also consider that the underlying *parameters* governing these events are random [11, 12]. The most simple method of achieving this is by applying a distribution to one or more of the key epidemiological parameters. For example, one may consider that the effective transmission rate, *β*, is drawn from some distribution. We can then find the solution to the ODE after having first drawn *β* from said distribution, assuming it then remains fixed. If we draw this random variable many times, we can again answer some probabilistic questions associated with the epidemic. In this case, we can consider the stochasticity to be *parameter-driven*. However, drawing a parameter from a distribution is still a simplification, and implies that there is uncertainty about a parameter that remains *fixed* over time; this has parallels to uncertainty propagation when estimating parameters. It is potentially more realistic to consider that the parameters themselves are stochastic processes; our underlying parameters are time-varying and noisy, as in [10]. However, this may still be unsatisfactory as the covariance (and autocorrelation) structure of the process may be more complicated than a simple noise term.

In this paper we consider the effects of setting a parameter to be the solution to a stochastic differential equation (SDE) - we term this parameter-driven randomness. Crucially, we also wish to maintain event-driven stochasticity. We refer to models in which there is both parameter-driven randomness and event-driven randomness as doubly stochastic. After first developing the methodological framework, we consider the impact of this exogenous stochasticity on the epidemic dynamics including the early growth, the risks of extinction and the long term dynamics especially the frequency of stochastic resonance [13].

## 2. Methodology

Throughout this paper, we shall be comparing our proposed double-stochastic model to the stochastic (CTMC) *susceptible-infected-recovered* (SIR) model which forms a basis for many models in the literature [14, 15, 16, 17]. We first introduce the deterministic SIR model with constant population size, governed by the following set of ordinary differential equations [14, 10]:

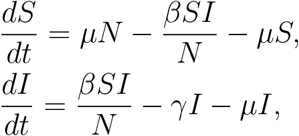

where *β* is the effective transmission rate, *γ* the recovery rate, *µ* is the birth / death rate and *N* is the population size (assumed constant). In this formulation *S, I* and *N* all refer to the number of individuals, making comparison with the CTMC models more intuitive.

The CTMC version can be thought of as the ‘natural’ extension of this system, where the transition rates in the ODE correspond to the rate at which events occur in a continuous-time Markov chain [7, 18]. In the continuous-time Markov chain formulation of the SIR model, the CTMC on the vector of states, **X** = (*S*_*t*_, *I*_*t*_), can be defined with transition rates from a state with *s* susceptibles and *i* infected individuals for a small time-step *δ*:

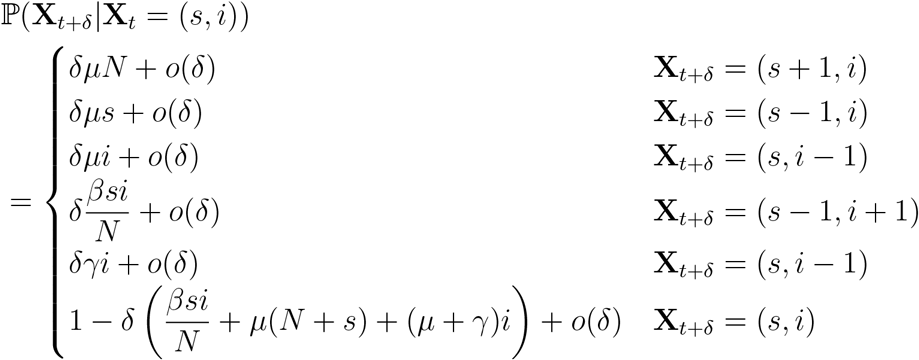

In models such as this, the parameters governing the transition rates are generally assumed to be constants, or occasionally deterministic functions of time. Examples of the latter are assuming *β* to be sinusoidal in models reflecting climatic forcing [19, 20], or for *β* to be a switch between high and low values representing increased mixing when children are in school (or some hybrid of these approaches) [21, 22].

### 2.1. The double-stochastic formulation

In this work, we shall consider that *β*_*t*_ is no longer constant, but instead is the solution to an Itô stochastic differential equation [23, 24, 25],

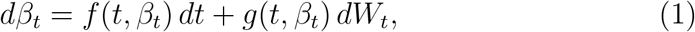

where *f* and *g* are the drift and diffusion functions, respectively, and *W*_*t*_ is a Wiener process. This gives rise to the ‘natural’ double-stochastic extension of the CTMC model above. We define the double-stochastic model as follows. Let 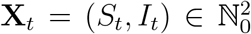 for all *t* ≥ 0 be a (time-inhomogeneous) stochastic process with transition probabilities,

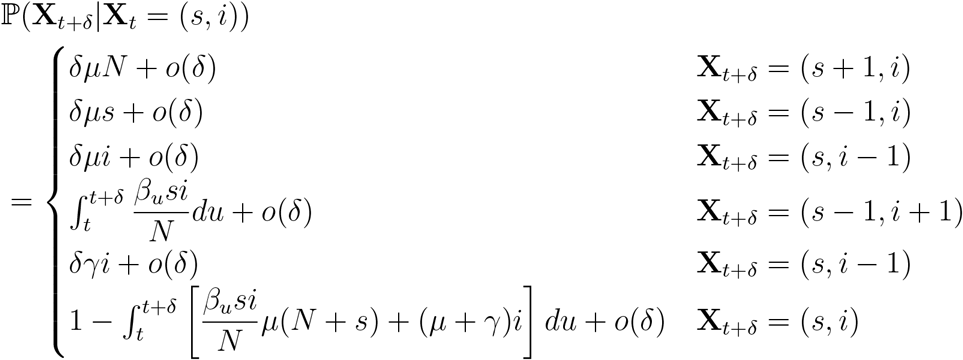

where *β*_*t*_ is the transmission rate at time *t*, given by the solution to the stochastic differential equation (equation 1), and the other parameters (assumed constant) are defined as in the standard model. Note that *β*_*t*_ is still a global parameter and so this model is still homogeneous across the population, but not in time. **X**_*t*_ is a double-stochastic SIR model with demography. Although we have introduced this additional stochasticity through the transmission rate, *β*_*t*_, other parameters could be made stochastic in an identical manner; however, we feel that noise in the transmission rate is the most epidemiologically natural assumption. Throughout this paper, we focus on only allowing *β* to be stochastic, as numerical simulations suggested that this has a more pronounced effect on epidemic trajectories than allowing *γ* to be stochastic. This formulation is very similar to the standard CTMC model, with the only difference being that terms involving *β* (*i*.*e. δβSI/N*) have been replaced with the time-integral 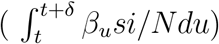.

We shall assume that the functions *f* and *g* in the SDE for *β* are selected such that *β*_*t*_ has a unique solution on the interval [0, *T*] for *T >* 0 (global Lipschitz continuity of *f* and *g* is a sufficient condition for this [24]). In addition, since *β* models a biological parameter, namely the effective transmission rate, we require that: (1) The solution 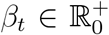 for all *t* ≥ 0; and (2) The solution is mean-reverting. That is, there is some known value which *β*_*t*_ is pulled towards, and the process will not stray too far from this value.

A suitable process is therefore the Cox-Ingersoll-Ross (CIR) model [26]. The solution to such an SDE exhibits the desired behaviours listed above, and is also well-studied in the finance literature, as it is often used in the modelling of interest-rate derivatives pricing [27]. It also has the advantage of having a known solution, and thus can be simulated exactly, without the need to employ approximate methods such as the Euler-Maruyama scheme [28].

### 2.2. The Cox-Ingersoll-Ross Model

The CIR model [26, 29] is the solution to:

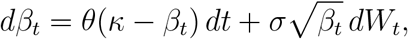

with parameters *θ, κ, σ >* 0.

For the purposes of epidemiological modeling, the Cox-Ingersoll-Ross model satisfies the requirements listed above - it is a non-negative and mean-reverting process. In this formulation, *θ* describes the strength of mean-reversion, *κ* informs the long-term mean value of the process and *σ* the volatility of the process. Furthermore, if 2*θκ* ≥ *σ*^2^ the process is strictly positive (otherwise it can occasionally touch zero) – this is known as the Feller condition. We shall assume throughout this paper that the Feller condition is met.

The Cox-Ingersoll-Ross model has the following two useful distributional properties. Firstly, it has a closed form solution. In particular, the process follows a non-central *χ*^2^ distribution,

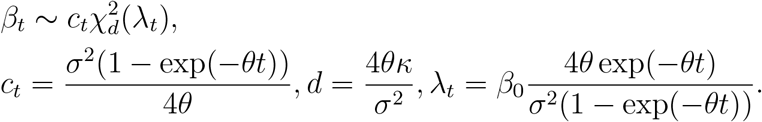

As a result, the process can be simulated exactly. Secondly, the distribution of *β*_*t*_ approaches a gamma distribution as time becomes large. In particular, the stationary distribution of the process is Gamma 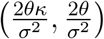 distributed. This is a much more approachable distribution and can be used for long-term calibration of the parameters of the SDE. More in-depth information on the properties of the CIR model can be found in section S1.

A natural question when using this model is the selection of the SDE parameter values (*θ, κ, σ*). In general, one could estimate values of *θ, κ* and *σ* from a discrete set of observations using, for example, ordinary least squares or maximum likelihood estimation [30, 31]. However, direct observations of *β* are unlikely (although climatic proxies are available if we knew the functional relationship); in principle it may also be possible to gain insights from estimates of the time-varying reproductive rate *R*_*t*_, but multiple challenges exist.

By modelling the parameters as CIR processes, we have made an implicit assumption that, over long timescales, our epidemiological parameter *β* will (approximately) have a Gamma distribution. We note that this is consistent with stochastic epidemiological models which assume a negative binomial distribution for secondary cases [32], as this arises from Poisson sampling with a Gamma distributed rate parameter. This long-term distribution helps with the selection of the SDE parameters, given the properties of the CIR model and Gamma distributions:

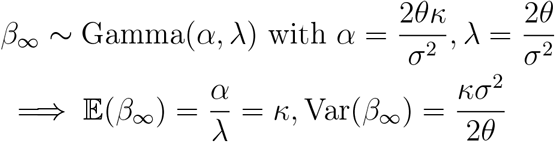

Generally, *κ* will be relatively straightforward to select when modelling *β*_*t*_ as it will be equal to the effective transmission rate in the standard case (i.e. the value of *β* used in the standard ODE or CTMC model). Having determined *κ*, the selection of *σ* and *θ* is less obvious. One method is to consider the required variance and half-life 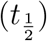 of the CIR process. For the CIR model, the half-life is given by 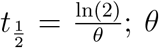 can then be selected so as to give the desired half-life, allowing *σ* to be determined from the variance (*σ*^2^ = 2*θ*Var*/κ*).

### 2.3. Simulation method

Here we briefly present a simulation method for the double-stochastic process. Efficient simulation is relatively straightforward for this method as the Cox-Ingersoll-Ross SDE has a known solution, hence the SDE can be simulated exactly by sampling from its distribution. We simulate the double-stochastic process by first finding a solution to *β*_*t*_ and then embedding this within the epidemic simulation using a *τ* -leap methodology [33]. The algorithm used for the simulation of the CIR model is presented in Algorithm 1.

#### Algorithm 1 Exact simulation of the CIR model

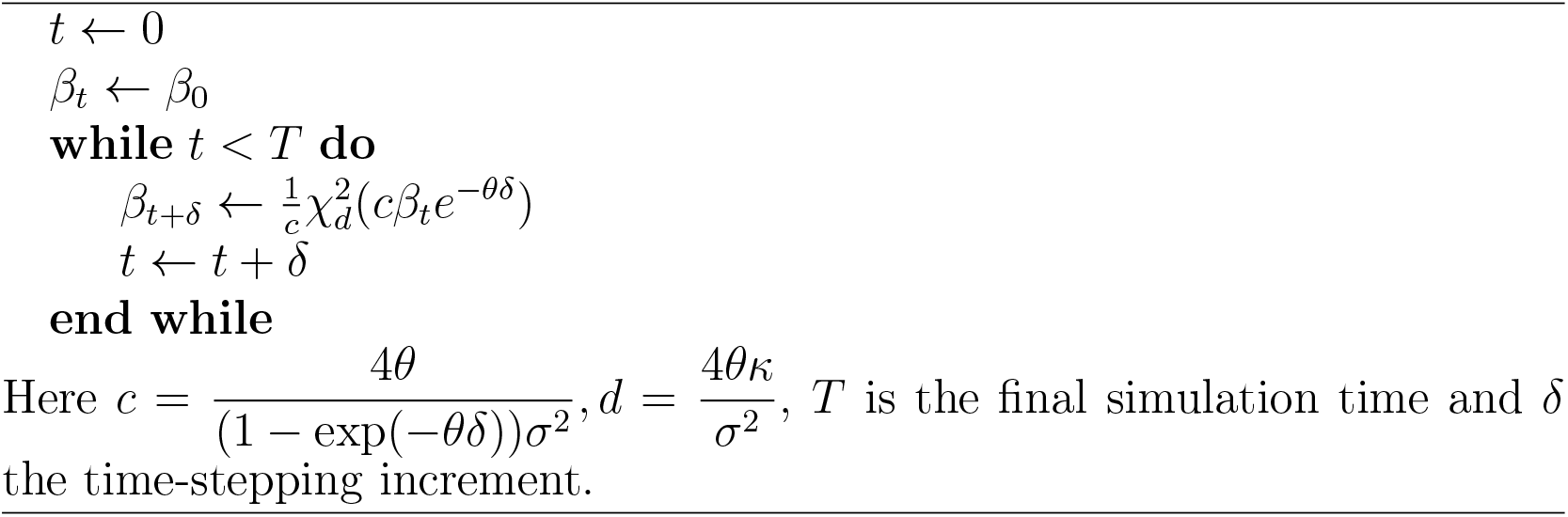

Once we have the values of *β*_*t*_ the epidemiological simulation method used throughout this paper is then the standard *τ* -leaping method [33] but with infection rates being 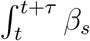 *ds* rather than *τβ* (where this integral is calculated numerically from the exact *β*_*s*_ values). This means that it is often useful to set *δ < τ* (so that steps in the SDE are smaller than steps in the epidemiological simulation), and to have *τ* as an integer multiple of *δ*, in order to get a more accurate value of this integral. Indeed in our simulations we take *τ* to be one day, but *δ* to be one tenth of this, as it was found that even for the most volatile process covered in this paper, the relative error at this granularity was approximately 1% (compared to the integral values found when *δ* was taken to be 1*/*10, 000 days).

If one wished to use a different SDE, then an exact simulation method is preferred if possible (that is, one that uses the distributional characteristics of the solution to calculate its simulated values, as in Algorithm 1). If a closed-form solution of the SDE is not possible, one instead must settle for approximate methods like Euler-Maruyama, stochastic Runge-Kutta or Milstein methods [34, 35]. Care must then be taken to ensure that these approximations are of sufficiently high order. For example, the Euler-Maruyama method is insufficient for finding solutions to the CIR model due to the non-linear diffusion term.

Since Algorithm 1 is an exact simulation method, it is very computationally efficient. Therefore, simulation of the double-stochastic process is comparable in terms of computation time to standard *τ* -leaping. While we do not present or use it here, there are methods for Gillespie simulation with a time-inhomogeneous effective transmission rate [36] (this works by taking an upper bound of the rate and then performing poisson-thinning). Unfortunately, this method does not work exactly with *β*_*t*_ being the solution to the CIR model as the process is unbounded. However, a threshold *β*^∗^ can still be selected such that the majority of the density of *β*_*t*_ lies below this density (and, as discussed, this density can be calculated exactly).

## 3. Results

We now present a range of theoretical and numerical results related to the double-stochastic model, and compare these to standard stochastic and deterministic models. For numerical results, we need to choose parameters to complete the simulations; here we pick two contrasting but epidemiologically relevant parameter sets: pertussis-like, and pandemic influenza-like parameters. The values for these regimes are shown in Table 1. It is important to note that this paper is *not* focused on highly accurate modelling of these particular diseases, but instead aims to present the method with plausible parameter values. These particular diseases allow us to do this – pertussis is slow yet highly infectious, and pandemic influenza is fast but much less infectious, and so these should serve to cover a wide range of modelling scenarios.

**Table 1:** Parameter values for different exemplar diseases. We note that the transmission rate *β* for the deterministic model (or *κ* in the double-stochastic model) is given by *β* ≈ *R*_0_*γ* – if small-scale demographic effects are ignored.

| Disease | $R_0$ | $\gamma$ (days) |
| --- | --- | --- |
| Pandemic influenza | 1.7 | 1/2 |
| Pertussis | 17 | 1/22 |

**Table 2:** Mean and variance values under the different stationary distributions for pan- demic influenza and pertussis.

|  | Pandemic influenza | Pertussis |
| --- | --- | --- |
| $\kappa$ value | 0.85 | 0.77 |
| Variance, high | 0.048 | 0.040 |
| Variance, medium | 0.016 | 0.013 |
| Variance, low | 0.010 | 0.008 |

Within these scenarios, we shall study a range of *θ, σ* pairs, corresponding to different variability and different rates of mean reversion. In particular, we shall focus on three underlying stationary distributions for *β*_*t*_ for each disease, relating to gamma distributions with different variances. We term these high, medium and low stationary variance.

### 3.1. Limiting behaviour in θ

Of the two parameters governing the stochastic dynamics of *β*, the strength of mean-reversion, *θ*, is potentially the one that will be most difficult to fit to available data (although we note that weather tends to have strong short-term correlations). Here, we therefore consider the limiting behaviour of the double-stochastic model when *θ* is either large or small.

We propose that, as *θ* → ∞ and the half-life is very short, **X**_*t*_ = (*S*_*t*_, *I*_*t*_) converges in the almost sure sense to the standard CTMC model with rate of infection *β* = *κ*. Whereas when *θ* → 0, provided that the stationary distribution of the process is held constant, *S*_*t*_, *I*_*t*_ converge in distribution to the standard CTMC model with rate of infection *β*_0_. That is, for fast mean-reversion, it is only the mean *β* that affects the dynamics; while for extremely slow mean-reversion the dynamics are dictated by the initial value of *β*. The proof of these statements can be found in section S2. These results make intuitive sense – when the strength of mean-reversion is very high, the epidemiological process averages across the very fast fluctuations and therefore experiences transmission rates close to the mean value, and when the strength is very low, the process hardly changes, and so will be almost constant at its initial value.

**Figure 1.**
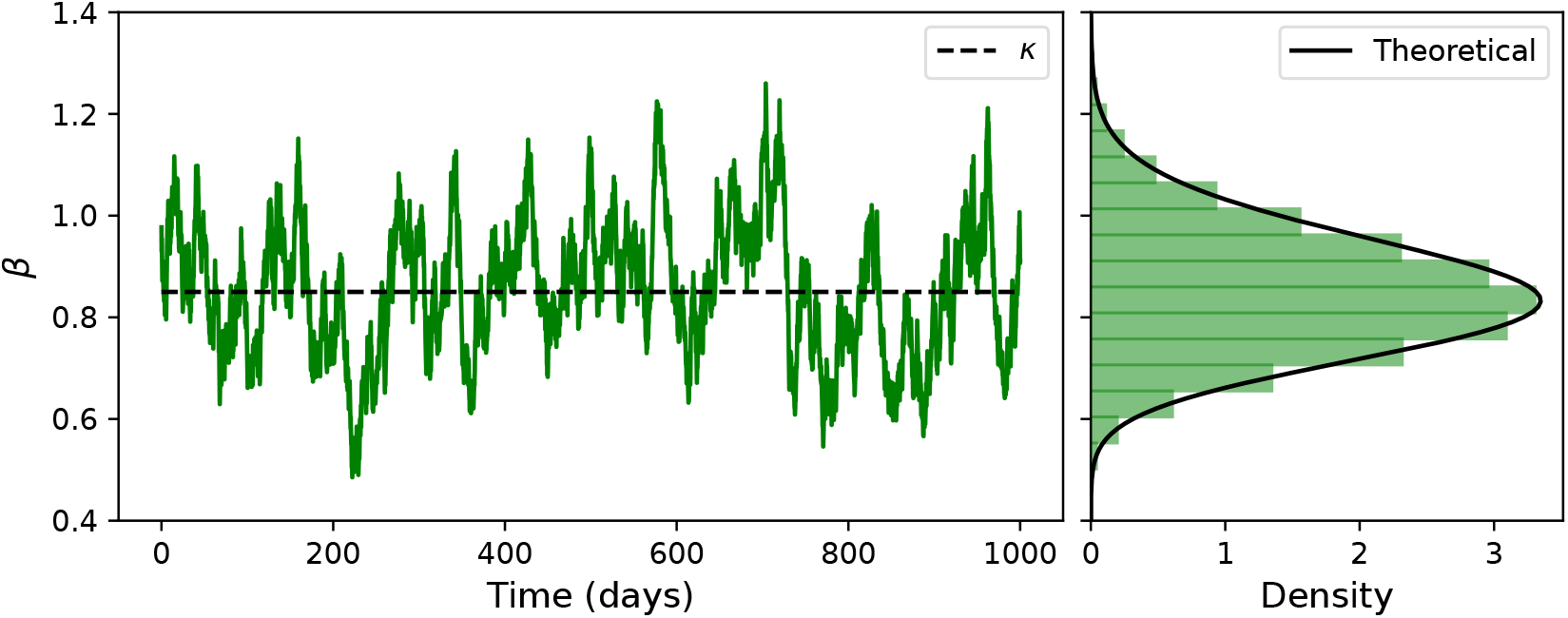
Left: sample trajectory of a Cox-Ingersoll-Ross SDE with *θ* = 0.07, *κ* = 0.85, *σ* = 0.05. Right: theoretical Gamma distribution in black with histogram of 1000 simulations of the SDE in green.

Figure 2 shows the means of epidemic trajectories with varying *θ*, but where the stationary distribution of *β* has been held fixed, reflecting the analytical results shown above. The figure also shows the two limiting cases. It is also worth noting that despite each simulation having the same stationary distribution, the trajectories vary significantly. Hence the mean outbreaks are not uniquely determined by the stationary distribution selection, the selection of *θ* is also key. As mentioned, we consider that selection of *θ* is generally a heuristic one, but it could also be selected based on exogenous variables for *β*_*t*_. For example, if we believe that air temperature is a key driver of *β*_*t*_, then we could consider that the parameters for *β*_*t*_ will be sim- ilar to those of an SDE model for air temperature. It has been estimated that a suitable value for the mean-reversion parameter of an SDE describing temperature is 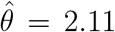, corresponding to a half-life of approximately 2.7 days [37].

**Figure 2.**
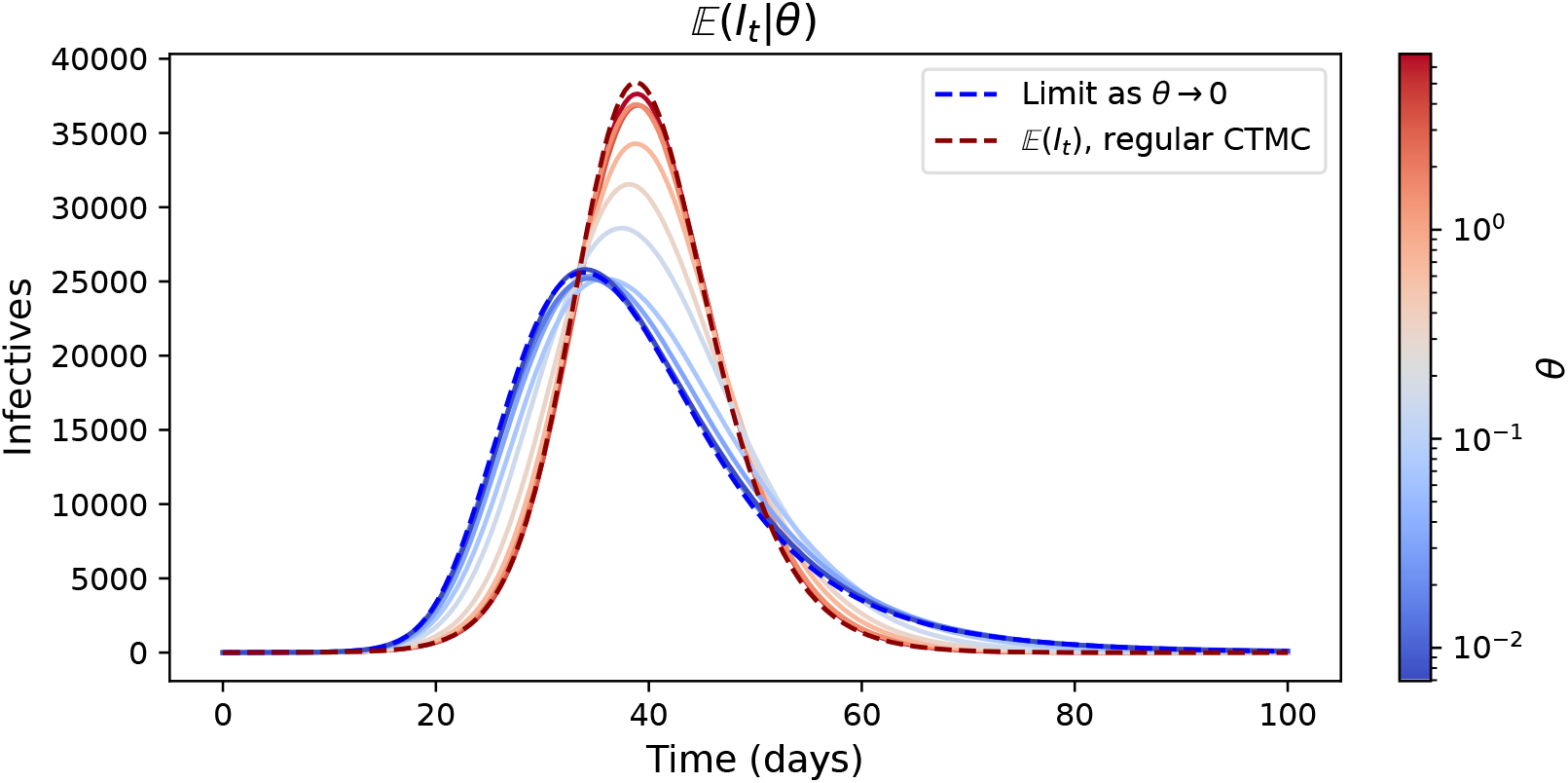
Mean of 1000 simulations under the Pertussis parameter regime with varying values of *θ*. Each simulation was initialised with a single infected individual in an otherwise entirely susceptible population with *N* = 10^6^. Note that in our simulations, *β*_0_ is drawn from the stationary distribution,0.85 and *σ* is scaled with *θ* such that *β*_*t*_ has the same stationary distribution in each simulation. Hence here the limit as the equivalently *θ* → 0 is the mean CTMC value when *β* is pre-drawn from the stationary Gamma distribution.

These limiting results show that in the limit of *θ* being either large or small, the sample trajectories are predictable and can be reduced to models within the scope of pre-existing models within the literature. This also provides an idea as to what happens when *θ* is close to these limiting results, and what we would expect to see on average as we move from a low *θ* to a high *θ*.

### 3.2. Early epidemic behaviour

Another important property of a model in epidemiology is the early behaviour of an outbreak. In particular, the early growth rate and the extinction probability (the likelihood of the disease not causing a large outbreak) are of utmost importance.

The early behaviour of an epidemic initialised close to the disease-free equilibrium can be studied through the use of a branching process (birth-death) approximation for the number of infected individuals in the continuous-time Markov chain system [10, 18]. This is justified by noting that close to the disease-free equilibrium (i.e. when the number of infected individuals is low), the infection term *βSI/N* ≈ *βNI/N* = *βI* and so new infections can be thought of as ‘births’ with rate *βI* and recoveries as ‘deaths’ with rate *γI*. This approach provides insights into both the growth rate and the extinction probability. To better conform to the branching process ideal, we shall consider *µ* = 0 since the early period of an outbreak is relatively short and so demographic events in the epidemiological model can be ignored.

#### 3.2.1. Early growth

One important metric is the growth rate of an outbreak in its early stages. Let *I*_*t*_ be the birth-death approximation of the *I* class, as discussed. Then, a standard result [38] is that:

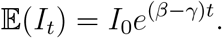

Now consider *I*_*t*_ to be the birth-death approximation of the double-stochastic SIR model with *β*_*t*_ the solution to the CIR SDE. In this case, we can use the result from the simple birth-death approximation by noting that the (*β* − *γ*)*t* term comes from integrating *β* − *γ* over time. Thus, we can simply replace the exponent by the time integral of *β*_*t*_ − *γ*,

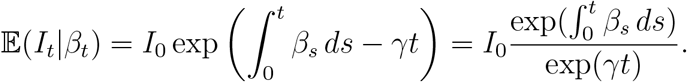

Then, by the tower property,

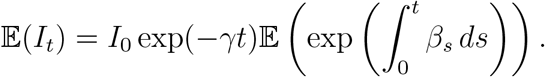

We note that, 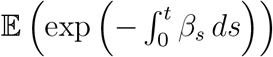 is the Laplace transform of the CIR model, and is is commonly used in mathematical finance for the price of a risk-free zero-coupon bond maturing in *t* years, where the interest rate is driven by a CIR process [26, 29]. As such, the solution to the expectation of the time-integral is a known result in the literature and is presented analytically in [26, 29]. Using this result gives that

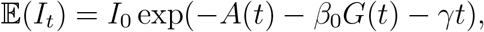

where

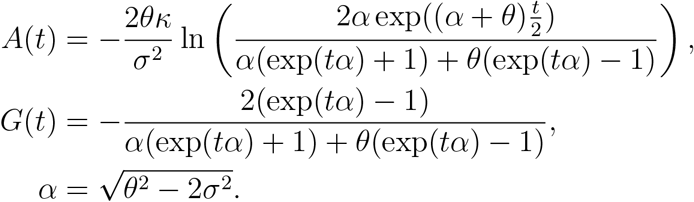

We can also study the asymptotic growth rate of E(*I*_*t*_) to more readily compare it to the standard CTMC model. That is we take time to be large, but still insist that the outbreak remains small relative to the population size. This gives,

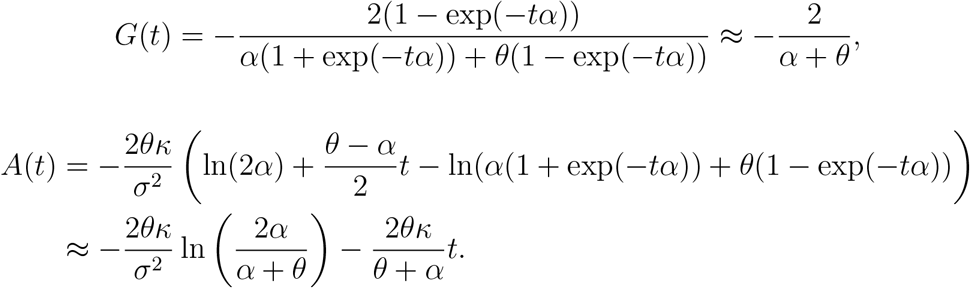

Combining these gives that, for large enough *t*,

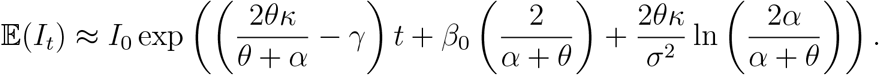

So, the asymptotic growth rate of E(*I*_*t*_) is,

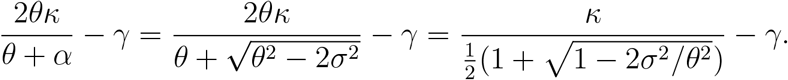

Which is strictly larger than the growth rate of the standard CTMC model (*κ* − *γ*). Intuitively, the additional stochasticity introduced by considering *β*_*t*_ to be stochastic increases the expected size of the *I*. Finally, note that the approximation given above may not be valid if 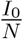 is not sufficiently small as susceptible depletion will occur prior to the asymptotic growth rate given in the approximation being reached.

#### 3.2.2. Early Extinction

Another important epidemiological statistic is the early extinction probability following the introduction of *i* infected individuals into a susceptible population. Again, we consider the birth-death approximation for the SIR model and its probability of extinction in finite time. For the standard CTMC SIR model, and for *i* ≥ 1 infected introductions, the early extinction probability is [39]:

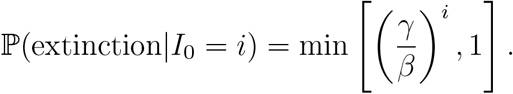

To study a result for the double-stochastic process, we shall also take a birth-death approximation of the process, where births occur at rate *β*_*t*_, and deaths at rate *γ*. Let 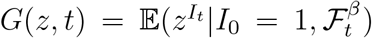 be the probability generating function of the process (conditioned on the filtration [24] of *β*). The forward PDE for *G* is [40],

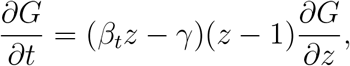

and can be solved using the method of characteristics, where

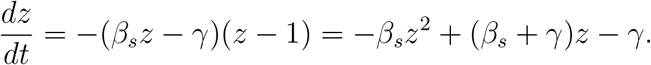

These are curves along which *G* is constant. In order to find the extinction probability *G*(0, *t*), we must consider a characteristic curve such that *z*(*t*) = 0, noting that since *G* is constant on this curve *G*(0, *t*) = *z*(0). We can solve the characteristic ODE by substituting *u*(*s*) = 1 − *z*(*s*), giving a Bernoulli equation,

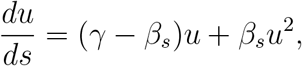

which yields the solution

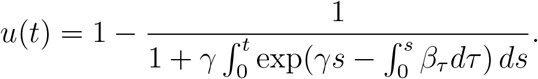

Therefore, we have that the extinction probability, conditional on the filtra- tion of *β* is,

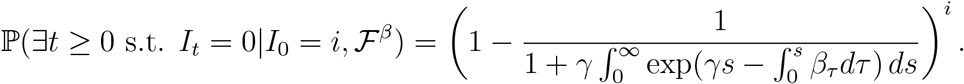

Then we can apply the law of total probability in order to find the unconditional extinction probability (i.e. not depending on the filtration):

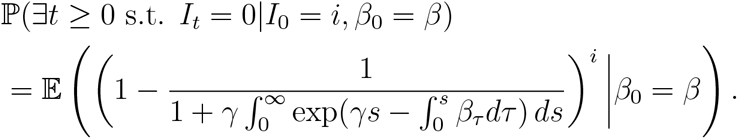

One difficulty is that, unlike in the standard case, the exponent *i* cannot be moved outside the expectation, because the lineages are no longer independent due to the presence of the correlated extrinsic noise in the form of the CIR model for *β*. Therefore, we shall instead attempt to form an equation allowing for the computation of this expectation. First, we expand the exponential term using the binomial theorem. Let 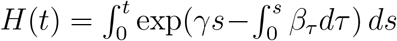, then

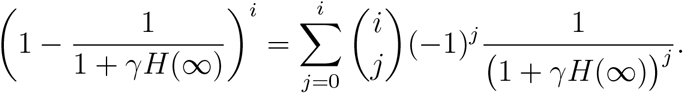

Note also that (via the so-called Schwinger parameterisation [41]),

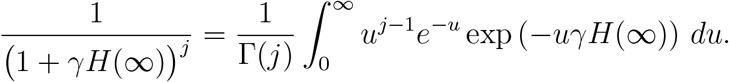

Thus, after taking expectations (and taking *λ* = *uγ* and Γ(0) = 1), it can be seen that the problem reduces to finding *m*(*β, λ*) = E exp(−*λH*(∞))|*β*_0_ = *β)*. Let *M*_*t*_ = E(exp(−*λH*(∞))|F_*t*_). Then, we may re-write this process as,

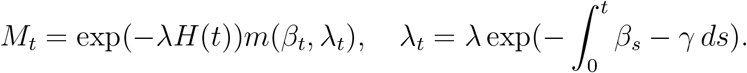

By noting that *M*_*t*_ is a function of *β*_*t*_, *λ*_*t*_ and *H*(*t*), we can apply Ito’s lemma to give

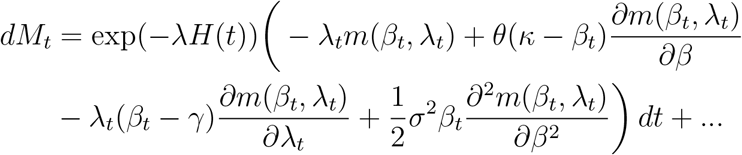

where we have omitted the diffusion term. In particular, *M*_*t*_ is a Doob- martingale and so cannot have a deterministic drift component (i.e. the drift term must be 0 at any time *t*). We can therefore extract the PDE from the drift term for *m*, in particular where we evaluate at *t* = 0, giving us the expectation term we desire, (*λ*_0_ = *λ, H*(0) = 0, *β*_0_ = *β*),

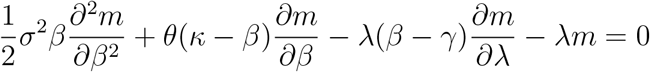

subject to the boundary condition *m*(*β*, 0) = 1. This PDE can then be solved for *m*, and substituted back into the formula for the extinction probability derived earlier.

Unsurprisingly, there is not an analytical solution to this PDE, and so the extinction probability must be computed numerically. Note that P(∃*t*≥ 0 s.t. *I*_*t*_ = 0|*I*_0_ = *i, β*_0_ = *β*) = 1 when *κ < γ*, as we would expect. This does not preclude there being an outbreak however, as the process *β*_*t*_, may have a period above 1, allowing an outbreak to take off.

We can form an analytical upper bound for the extinction probability using Jensen’s inequality for

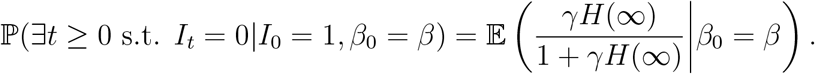

Since *γx/*(1 + *γx*) is a concave function, so Jensen’s inequality gives that

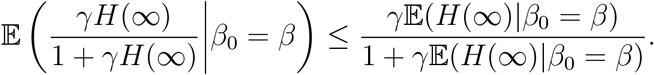

We can evaluate the integral,

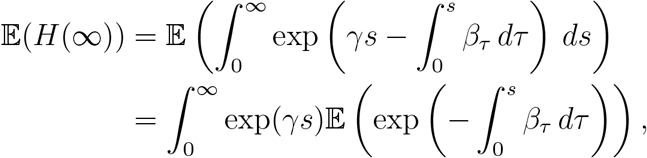

where the expectation is again the Laplace transform of the CIR model, as noted in the previous section. Thus, we have that,

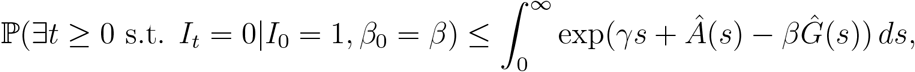

where

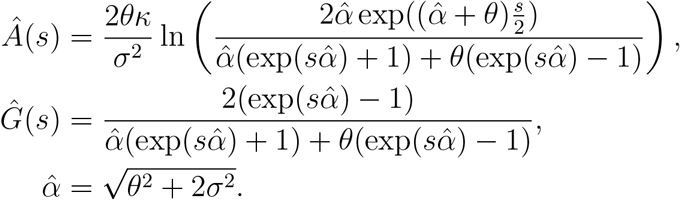

We show the extinction probability in the pandemic influenza and pertussis parameter regimes in Figure 3. In this figure, we see that in regions where the strength of mean-reversion, *θ*, is large, the probability of extinction is approximately that of the CTMC case (this is unsurprising given the result we proved in subsection 3.1). Similarly, when *θ* is small, the half-life of the process is much larger than the average infectious period, and so during the first and second infections in the outbreak (which account for the majority of the extinction probability), *β*_*t*_ will not change significantly, and so again, since *β*_0_ = *κ* in our simulations, the extinction probability in this region is approximately that of the CTMC case. The only region in which the extinction probability differs markedly from that of the base case is when the strength of mean-reversion is moderate — it is not so big that the process fluctuates frequently enough such that the effective transmission rate is approximately *κ*, but is larger enough so that *β*_*t*_ is still relatively volatile.

**Figure 3.**
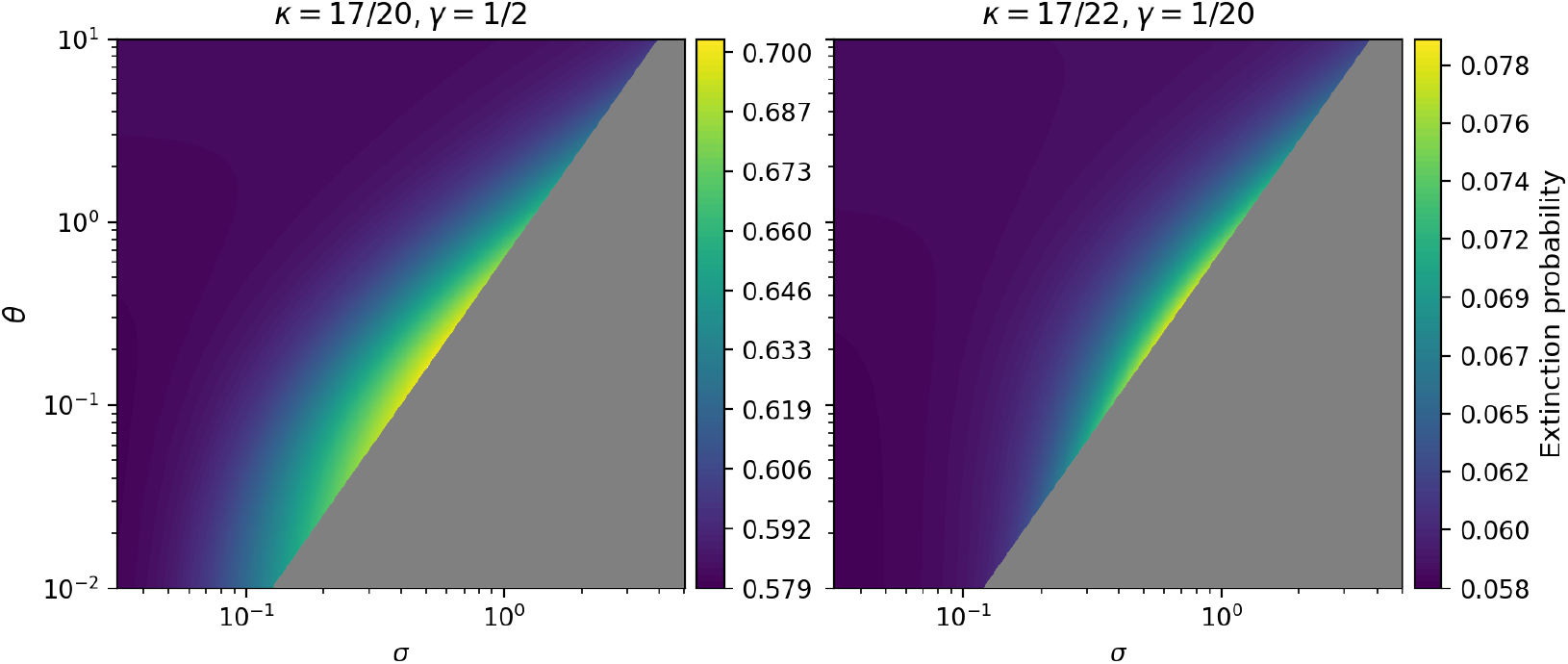
Heatmaps showing extinction probability with *I*_0_ = 1 under the pandemic influenza and pertussis (*κ, γ* pairs), with *β*_0_ = *κ*. Dark-grey regions indicate areas in which the Feller condition does not hold.

The birth-death approximation only holds when *S* ≈ *N*, and thus we also present numerical results showing how the extinction probability varies later in the epidemic, when this condition on *I* no longer holds. We maintain the assumption that *µ* = 0, such that there are no demographic processes, hence the infection inevitably goes extinct over long time scales. We therefore compute the probability that the infection is extinct at time *t* (Figure 4). We expect this to start at zero, plateau at the theoretical branching process value over epidemic time scales and then reach one once we expect the outbreaks to be over. For the pandemic influenza parameters (Figure 4, upper panels), inducing stochasticity on the rate of infection in this manner produces results which ‘smooth out’ the extinction probability, and that this effect becomes more extreme as the variance and the rate of mean-reversion of the CIR process is decreased. When realisations of *β* experience initial growth above the mean and stay at this elevated level, the population experiences epidemic burnout faster, and so (given that the outbreak takes off), the extinction time is earlier in these instances. The opposite is true for outbreaks with later extinction times. When *θ* is low and *σ* is high, these more extreme trajectories of *β* are most likely, as can be seen in Figure 4. Therefore although the branching process extinction probably is higher for high variance *β* (dark blue line), as seen in the initial plateau, there are later times when the extinction risk is less. Under the pertussis parameter regime however, there is very little difference between any of the empirical extinction probabilities plotted (Figure 4, lower panels). This is because, when initialised from close to the disease-free equilibrium, the much higher *R*_0_ of pertussis causes the deterministic forcing effect to be much stronger than in the case of pandemic influenza. Since the stochasticity of *β* is much smaller for pertussis relative to its *R*_0_ value, changes in the parameters of *β*_*t*_ make very little difference to outbreak trajectories.

**Figure 4.**
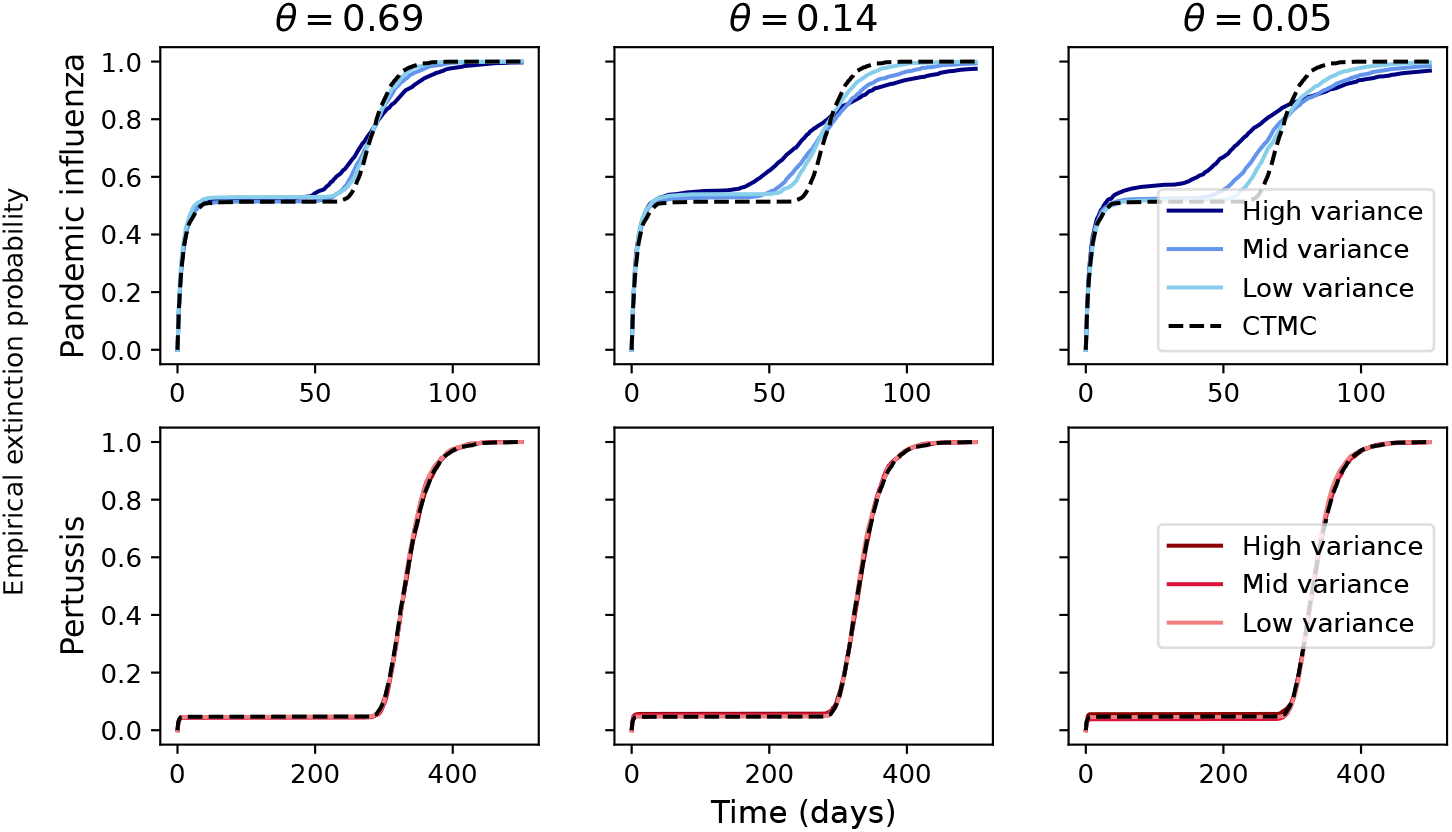
Empirical extinction probability under the pertussis (lower panels) and pandemic influenza (upper panels) parameter regimes. Each probability was produced from 2000 simulations, each initialised with one infected individual in an otherwise entirely susceptible population, with *µ* = 0 and *N* = 10^5^.

### 3.3. Long-term epidemic behaviour

The long-term dynamics of epidemiological systems are also of interest. In 1956, Bartlett [6] noted that in SIR-type models with demography and event-driven stochasticity, levels of infection will fluctuate about the endemic equilibrium level in a phenomenon called stochastic resonance (this differs to deterministic models where the oscillations are damped over time). We shall discuss how these fluctuations differ in the double-stochastic case to the standard CTMC model by comparing the power spectral densities. The power spectral density (PSD) of the infected class measures the intensity of every frequency of oscillation (essentially describing how often an increase in infection above baseline is likely to occur).

Traditionally to find the power spectral density, one approximates the CTMC as a linear SDE using the van Kampen system size expansion [42, 43]. We shall instead use the SDE approximation method in [18] for ease (though it gives the same underlying SDE), which via similar mathematical reasoning to the system size expansion, for our double-stochastic process yields

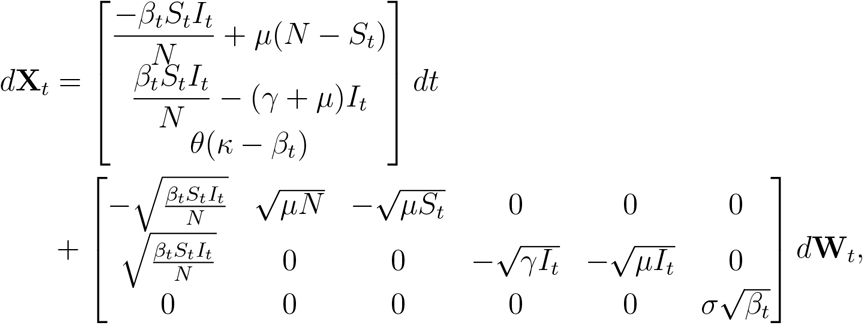

where 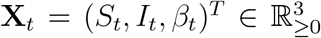. We then linearise this SDE about the en- demic equilibrium, 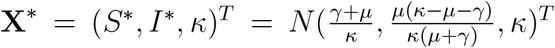. This linear approximation is

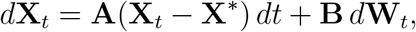

with

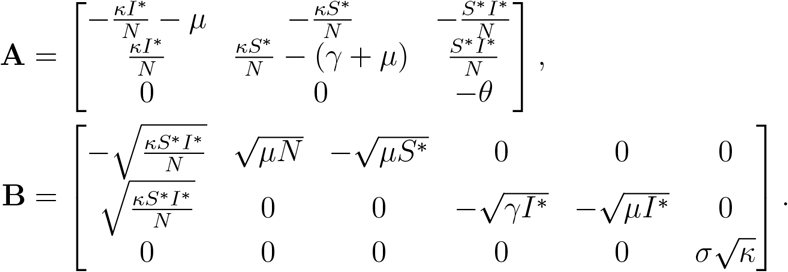

In particular, this approximation is an Ornstein-Uhlenbeck (OU) process [44]. For SDEs of this form, the Power Spectral Density, **S**(*ω*), is [44]

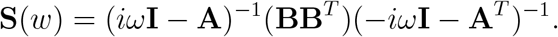

This gives an analytical approximation for the power spectral density of the double-stochastic process. The fully expanded version of this approximation is very long and not in itself informative and therefore is not written here, we instead present a numerical computation of the approximation (Figure 5).

**Figure 5.**
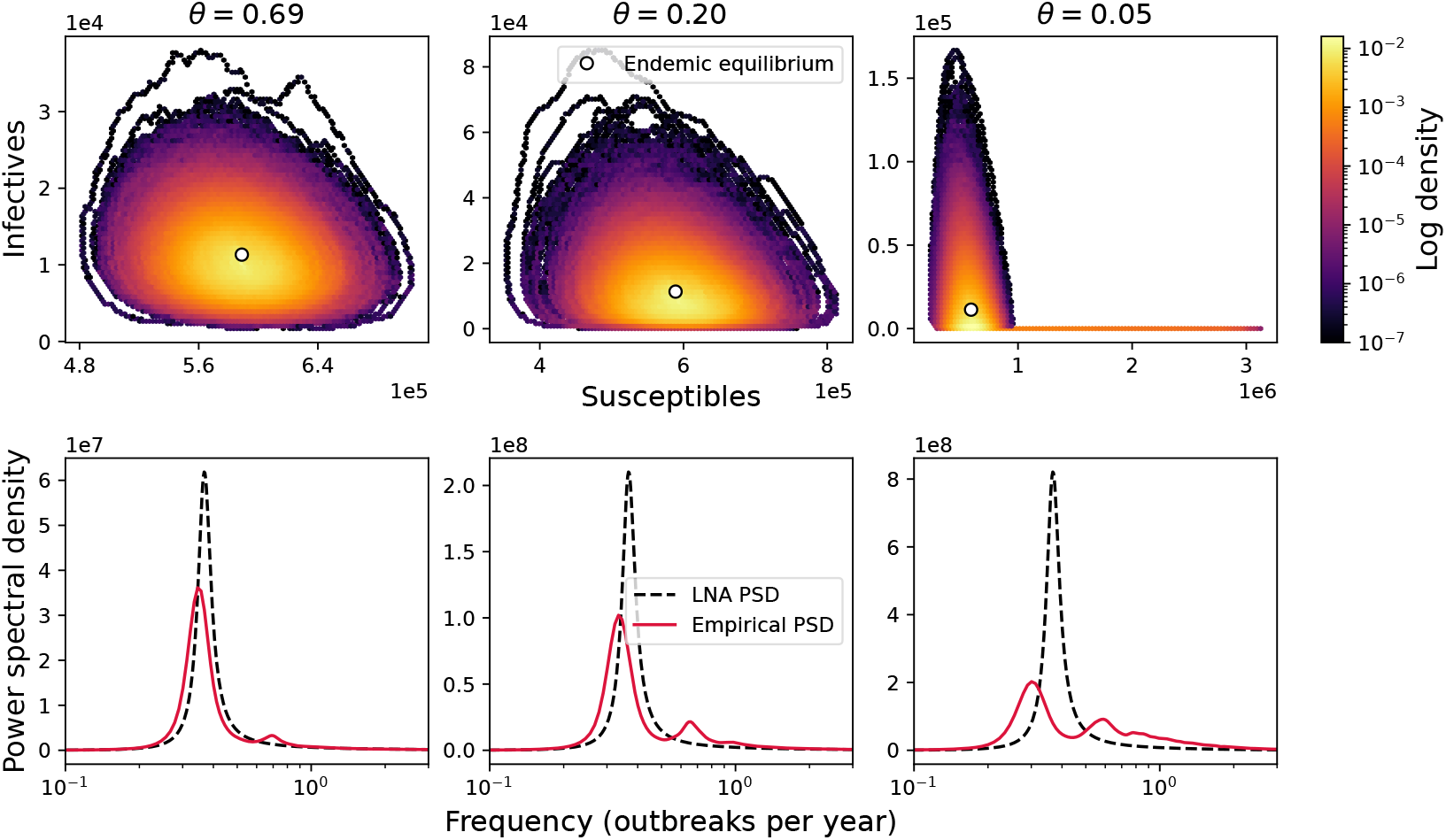
Top: empirical density of (*S, I*) taken over 1000 simulations under the pertussis parameter regime and each *θ* value. Each simulation was run for 10000 days at a time interval of 1 day. Bottom: theoretical PSD as found using the linear noise approximation outlined above and empirical PSD, taken by averaging the Fourier transform of each of the 1000 simulations. In each plot, the stationary distribution of *β* is identical.

We observe that the peak power spectral density approximation derived via the system-size expansion works well when there is rapid mean-reversion, but is less accurate when *θ* is lower. The reason for this is two-fold: in the full SIR model, the non-linear deterministic forcing prevents any class from being negative; and in models with event-driven stochasticity, the process may go extinct. Both of these are signatures of non-linear behaviour far from the fixed point, that the van Kampen approximation does not capture.

Indeed, since the approximation is an OU process, the distribution of the SDE is Gaussian. The mean is simply the steady state about which we linearised the system, and the covariance matrix, **Σ**, can be calculated via the continuous-time Lyapunov equation,

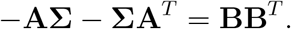

So, the stationary distribution of our linear approximation is Gaussian with mean **X**^∗^ and variance-covariance, **Σ**. This approximation is thus an ana- logue for the approximation used in [43], but has a different form due to the stochasticity in *β*.

While the Gaussian approximation allows for analytical results, and, as mentioned, is analogous to methods used to study more simple models, it also causes some problems. Since Gaussians exist in ℝ^*n*^, a portion of the density function will be negative. When this portion is small, we do not expect this to cause significant errors, but for certain (*θ, σ*) pairs, a large portion of the marginal density of the infected (*I*) class falls below 0. When this is the case, the approximation begins to break down (values of *I* below zero are not feasible in the stochastic model and are biologically meaningless). In Figure 6, we show the amount of density below 0 in the *I* class under the pertussis parameter regime for a range of *θ* and *σ* values. Our linearised approximation will be poor in areas where this probability is higher. Indeed, this is what occurs in Figure 5. The left plot has a half-life of 1 day, equivalently *θ* ≈ 0.69 and the right plot *θ* ≈ 0.05, and we see in Figure 6 that this will increase the proportion of the density that is negative in the OU approximation.

**Figure 6.**
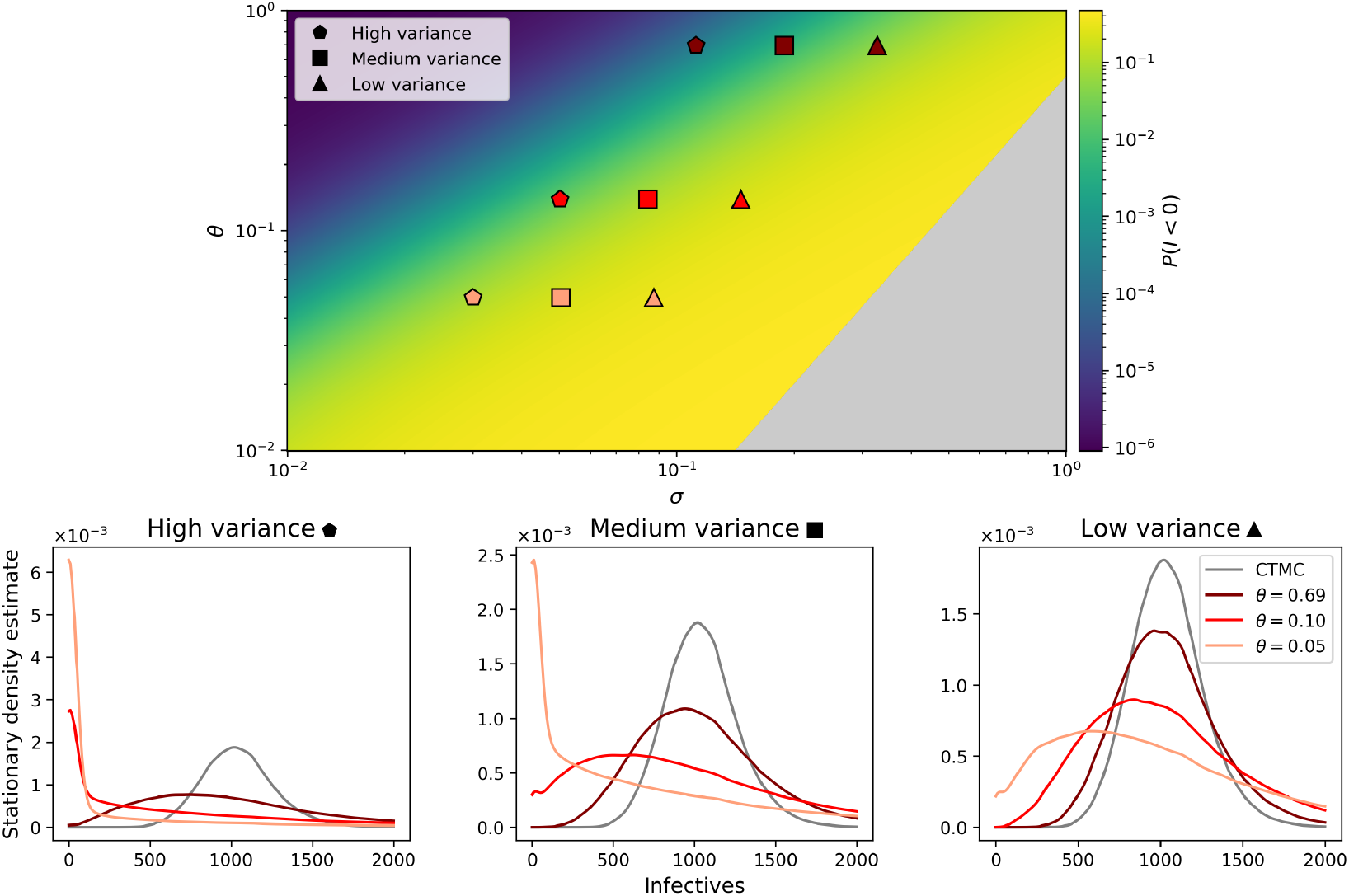
Stationary kernel density estimates of the double-stochastic process under the pertussis parameter regime from 10000 realisations initialised at endemic equilibrium with *N* = 10^6^ over 25 years.

This is reflected in stationary density estimates presented in Figure 6. We see that for sufficiently small mean-reversion half-lives (and particularly in the CTMC limit) the stationary density of infecteds is roughly Gaussian, but this begins to break down as the half-life increases (or equivalently as the strength of mean reversion decreases), this is particularly pronounced as the variance of the stationary distribution increases. The high-variance realisations with the weakest mean-reversion (*θ* = 0.05) is the most extreme example of this, where a significant amount of the density is at *I* = 0, i.e. that the disease goes extinct.

In terms of trajectories, what occurs here is that when initialised close to endemic equilibrium, we still observe resonance behaviour, as we would expect, but outbreak trajectories become more ‘spiky’ when *σ* or *θ* are increased (Figure 7). That is, with higher stationary variance and weaker mean-reversion, stochastic outbreaks are often significantly larger than in the standard continuous-time Markov chain model (as there are prolonged periods with high *β* values), and between these outbreaks the level of infection may be significantly smaller. In keeping with the standard CTMC version of the SIR model, when the system is close to the endemic equilibrium, the stochastic effects are significantly stronger relative to the deterministic forcing (as the deterministic rates of change are close to zero); the opposite is true further from the endemic equilibrium and in particular close to the disease-free equilibrium. However, in both scenarios, large *σ* or small *θ* leads to far greater stochastic effects in our double-stochastic model compared to the CTMC.

**Figure 7.**
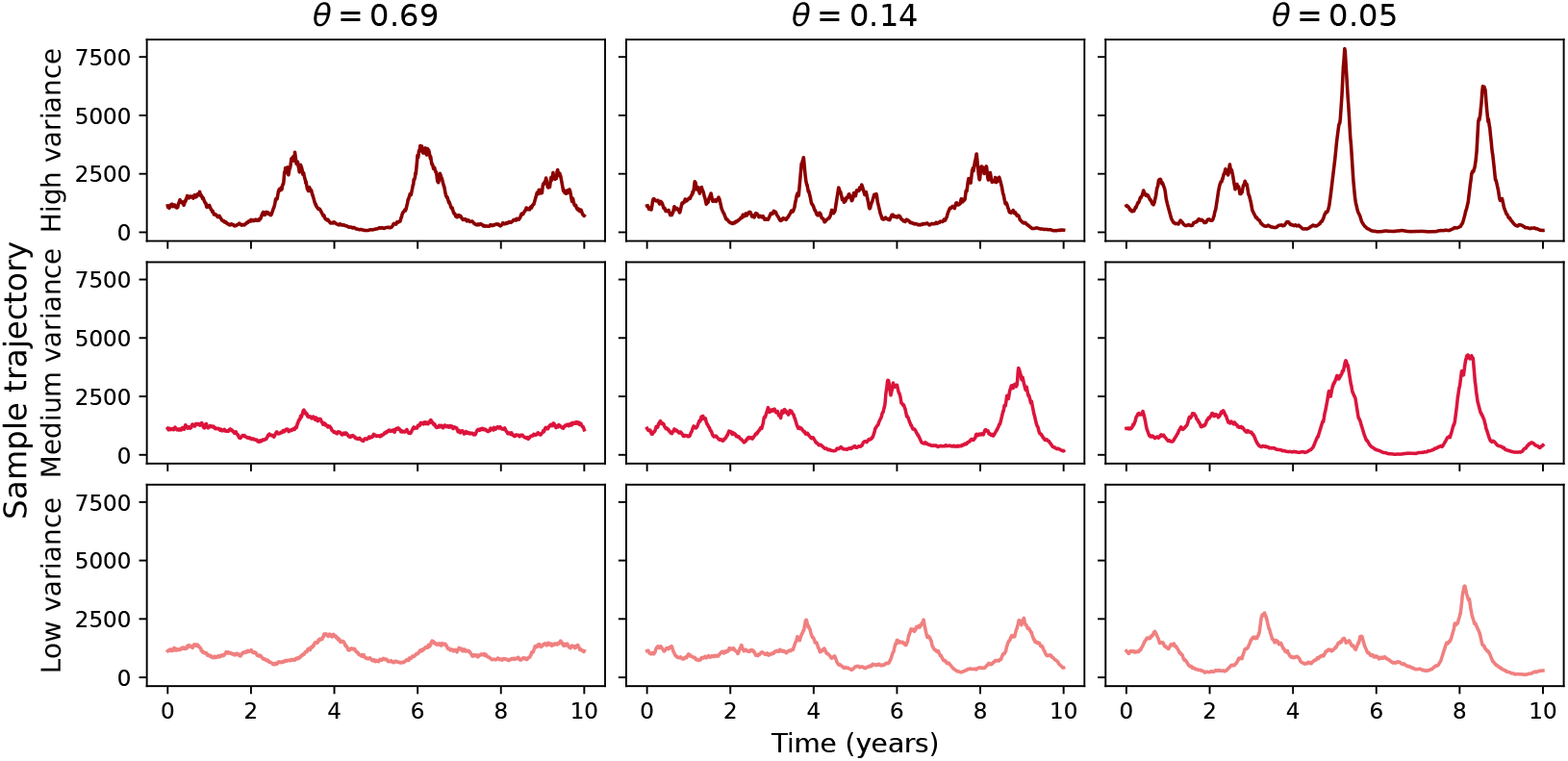
Sample trajectories of the double-stochastic process under the pertussis parameter regime initialised from endemic equilibrium with *N* = 10^6^. From left to right, half-life increases (and thus strength of mean reversion, *θ* decreases). From top to bottom, variance of the stationary distribution decreases.

In summary, the introduction of parameter-driven stochasticity into the SIR model in the form of an SDE for *β* generally results in more variation in outbreak trajectories, which is perhaps unsurprising. In the long-term, extinction is strictly more likely to occur in the double-stochastic model than when no parameter forcing is present. When the system is close to endemic equilibrium, more ‘peaky’ outbreak trajectories are seen, when compared to the standard CTMC model. In either case, smaller values of *θ* (equivalent to larger half-lives of *β*) and larger values of *σ* lead to the most extreme results – leading to a higher extinction probability, larger outbreaks and the biggest deviation from the standard models in the literature.

## 4. Discussion

We have presented an epidemiological model, novel for its inclusion of stochastic effects both in events and in the transmission parameter. We specifically focused on the SIR model with demography due to its simplicity and frequent use, but the methods presented can be applied to allow arbitrary compartmental models to be extended in such a manner, and for alternative stochastic parameters. For this model type with the Cox-Ingersoll-Ross SDE for the rate of infection parameter, analytical results were presented, where possible, for both early behaviour and long-term behaviour of an outbreak.

We showed that in the limit as *θ*→ ∞ (fast mean-reversion of *β*), the double stochastic process converges almost surely to the standard CTMC model from which it is adapted. Similarly, as *θ* → 0, the double stochastic process converges almost surely to a standard CTMC with effective transmission rate *β*_0_. These convergence properties give analytical explanations for our observed numerical results, and highlight the link between our double-stochastic results and the standard models present in the literature.

We also studied the early behaviour of the double-stochastic model, leading to an analytical expression for the early growth of infection. While in very early stages of an outbreak, growth maybe larger or smaller than in the CTMC case, in the large-time limit, regardless of the values of *θ* and *σ*, the growth-rate is higher than in this standard case. Furthermore, using the birth-death approximation, we found a method of calculating extinction probability, as well as an analytical upper bound on this quantity.

Results for long-term behaviour about endemic equilibrium were also considered. Under some parameter regimes, the double-stochastic process near the endemic equilibrium is well approximated by an Ornstein-Uhlenbeck SDE. The Gaussian nature of this approximation makes analysis significantly easier, for example giving rise to an analytical form of the power spectrum density. We discussed when this approximation holds and when it begins to break down. In particular, larger values of *σ* and smaller values of *θ* (increasing the conditional variance of the *β* process) reduce the validity of the OU approximation. Numerically, we saw that this is because under these conditions, outbreak trajectories become more extreme, yielding much larger outbreaks, followed by near-extinction.

The importance of stochasticity in epidemiological modelling has long been recognised, but the assumption of the standard continuous-time Markov chain models that biological parameters are fixed may be unreasonable. We have shown that relaxing this assumption leads to models which are well-behaved, but generate different results from the CTMC models often used in the literature. Indeed, the double-stochastic model enables a richer variety of outbreak dynamics, and thus potentially increased accuracy to real-world data.

Double-stochastic models of this type may be particularly useful when seeking to account for exogenous factors which are stochastic in nature (an example of such a variable is temperature). One obvious extension therefore is to consider models which have both deterministic and stochastic seasonality. This may be done through the use of a sinusoidal function in the drift term of the CIR SDE to replace *κ*:

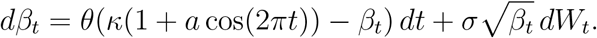

One difficulty with more complex models such as this (or indeed a large number of other SDEs which may be selected instead of the CIR model) is that the analytical results discussed within this paper may either become significantly more challenging to calculate, or intractable as the SDE may not have a closed-form solution. Furthermore, the numerical simulation of such a model becomes tougher as exact simulation methods cannot be exploited if such a solution is not known.

While the methods presented in this paper can be applied to models with more structure (for example susceptible-exposed-infected-recovered models), the analysis done is specifically on an SIR-type model which may be overly simple for some applications. In addition, while the CIR model has desirable properties as discussed, it is not bounded, and therefore it is possible that *β* becomes very large. This can be avoided with high probability with suitable selection of parameters, and using the CIR model avoids having to use an SDE without an analytical solution. Indeed, if one wished, parameters could be selected such that the vast majority of the density of *β* were to lie within the maximum and minimum of a sinusoidal seasonal model for *β*.

As this work has presented a new approach to modelling of outbreaks, a natural next step is to examine in more detail questions related to parameter selection and model fitting. Additional work is required to give modellers the full suite of tools needed to use the double-stochastic model for real-world applications. However, the results presented above demonstrate the potential of the double-stochastic model as an effective and analytically tractable approach.

## CRediT authorship contribution statement

DRH: Writing – review and editing, Writing – original draft, Visualization, Project administration, Methodology, Investigation, Formal analysis, Conceptualization.

MJK: Writing – review and editing, Supervision, Methodology, Investigation, Conceptualization, Funding acquisition, Conceptualization.

LD: Writing – review and editing, Supervision, Methodology, Investigation, Conceptualization, Funding acquisition, Conceptualization.

## Acknowledgements

DRH, MJK and LD were supported by the Engineering and Physical Sciences Research Council through the MathSys CDT (grant number EP/S022244/1). MJK was also supported by the Wellcome Trust (226057/Z/22/Z).

## Supplementary material

## S1. Methods

### The Cox-Ingersoll-Ross model

The CIR model has the following properties [29],

1. The conditional expectation is E(*β*_*t*_|*β*_*s*_) = *κ* + (*β*_*s*_ − *κ*)*e*^−*θ*(*t*−*s*)^.
2. As a result, *β*_*t*_ is mean reverting. That is, lim_*t*→∞_ E(*β*_*t*_|*β*_*s*_) = *κ* for any *β*_*s*_.
3. The conditional variance is Var 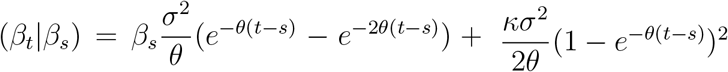
4. Therefore, *θ* can be thought of as the strength of mean-reversion, *κ* the long-term mean and *σ* the volatility of *β*_*t*_.
5. The distribution of the CIR process can be computed. In particular, we have that

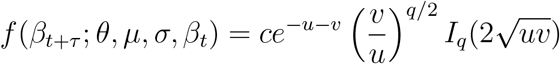

where 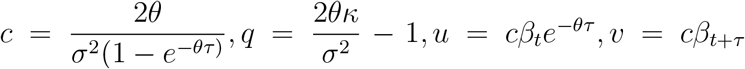 and 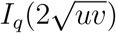 is a modified Bessel function of the first kind of order *q*. That is, it follows a non-central *χ*^2^ distribution.
6. The distribution of *β*_*t*_ approaches a gamma distribution as time becomes large. In particular, 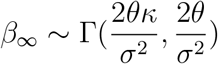.
7. When the Feller condition is met, that is if 2*θκ* ≥ *σ*^2^ and *X*_0_ *>* 0, then *X*_*t*_ is strictly positive; otherwise it can occasionally touch zero.

#### S1.1. Numerical integration of the CIR model

**Figure S1.**
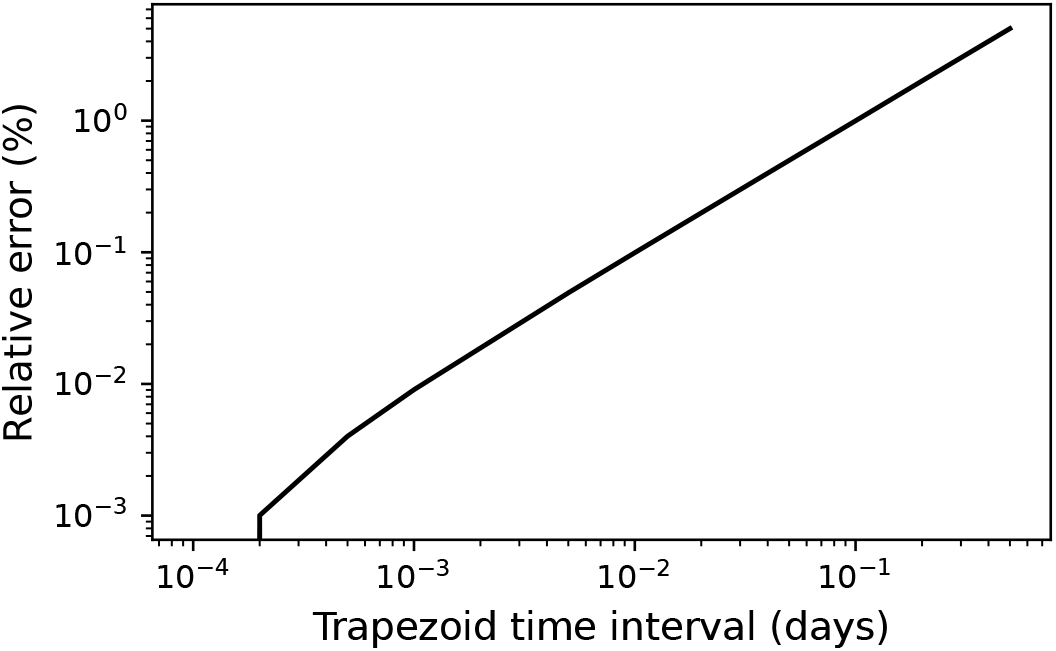
Relative error of trapezoid method for numerical integration of the CIR model over time, compared to with a time interval of 1*/*10000 days.

## S2. Results

### Proof of limiting behaviour in θ

We wish to show that the double stochastic process with mean-reversion parameter *θ* → ∞ converges in distribution to the standard CTMC case with effective transmission rate *κ* when *θ*. Also, that as *θ* → 0 (with the stationary distribution held constant), the double stochastic process converges in the almost sure sense to the standard CTMC case with transition rate *β*_0_. We first note that converge in *L*^1^ and convergence in *L*^2^ both imply convergence in probability, as a result of the Markov inequality.

We shall first show that

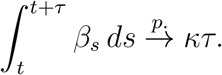

The solution to the CIR SDE can be used to find an expression for the time integral,

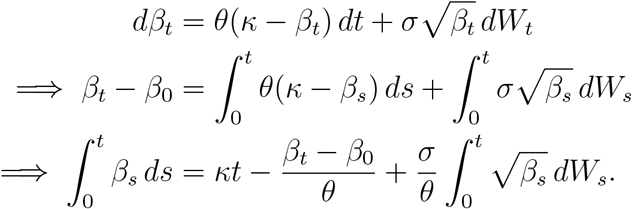

We shall study this term-by-term. Firstly,

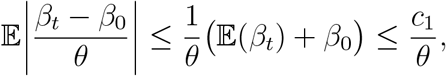

for some constant *c*_1_. The second inequality follows since E(*β*_*t*_) is bounded (as can be seen in section S1). As a result, this term exhibits *L*^1^ convergence to 0. As mentioned previously, therefore the term converges in probability to 0.

We now look at the integral (martingale) part,

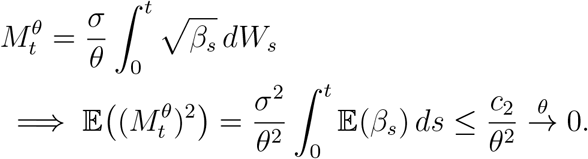

In other words, the martingale term converges in *L*^2^ to 0, and so converges in probability to 0. Combining these terms gives that,

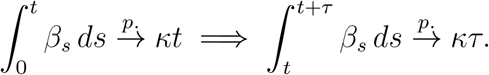

Define 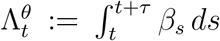. Then, there exists a continuous function *f* that maps 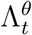 to the transition probabilities of the double-stochastic process. By the continuous mapping theorem,

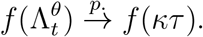

Then, noting that convergence in probability implies convergence in distribution, we appeal to the Portmanteau theorem. In particular, we shall use that for a sequence of random variables *X*_*n*_,

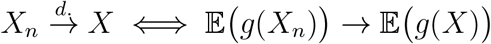

for every bounded continuous *g*. Therefore (noting that 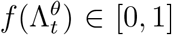 as it is a probability) we may apply the Portmanteau theorem on *g*(*x*) = *x*,

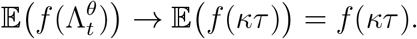

Then, since 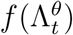 can be considered as being the transition probability of the double-stochastic process under a given path of *β* (i.e it is conditioned on 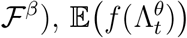 is the transition probability of the unconditioned process (by Tower property). As a result,

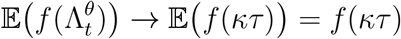

implies the result.

For the second statement, in order for the process to retain the same stationary distribution, *σ* must be scaled such that it remains proportional to 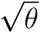.

Clearly, as *θ* → 0 both the drift and diffusion tend to 0, and thus the process does not move from *β*_0_. It follows that the integral tends to *tβ*_0_.

